# Tellurium: A Python Based Modeling and Reproducibility Platform for Systems Biology

**DOI:** 10.1101/054601

**Authors:** Kiri Choi, J. Kyle Medley, Caroline Cannistra, Matthias König, Lucian Smith, Kaylene Stocking, Herbert M. Sauro

**Author notes:** These authors contributed equally to this work (Kiri Choi, J. Kyle Medley).

## Abstract

In this article, we present Tellurium, a powerful Python-based integrated environment designed for model building, analysis, simulation and reproducibility in systems and synthetic biology. Tellurium is a modular, cross-platform, and open-source integrated development environment (IDE) composed of multiple libraries, plugins, and specialized modules and methods. Tellurium ensures exchangeability and reproducibility of computational models by supporting SBML (Systems Biology Markup Language), SED-ML (Simulation Experiment Description Markup Language), the COMBINE archive, and SBOL (Synthetic Biology Open Language). Tellurium is a self-contained modeling platform which comes with a fully configured Python distribution independent of other local Python installations on the target machine. The main interface is based on the Spyder IDE which has a highly accessible user interface akin to MATLAB (https://www.mathworks.com/). Tellurium uses libRoadRunner as the default SBML simulation engine due to its superior performance, scalability and ease of integration. libRoadRunner supports deterministic simulations, stochastic simulations and steady state analyses. Tellurium also includes Antimony, a human-readable model definition language which can be converted to and from SBML. Other standard Python scientific libraries such as NumPy, SciPy, and matplotlib are included by default. Additionally, we include several user-friendly plugins and advanced modules for a wide-variety of applications, ranging from visualization tools to complex algorithms for bifurcation analysis and multi-dimensional parameter scanning. By combining multiple libraries, plugins, and modules into a single package, Tellurium provides a unified but extensible solution for biological modeling and simulation.

## 1 Introduction

The field of systems biology is replete with software tools designed to assist in creating and simulating biochemical networks (Sauro and Bergmann, 2009). Examples of popular tools include COPASI (Hoops et al., 2006), CellDesigner (Funahashi et al., 2003; Funahashi et al., 2008), iBiosim (C.J. Myers et al., 2009), JigCell (Vass et al., 2006). PathwayDesigner (formerly JDesigner, Sauro, 2001), PySCeS (Olivier et al., 2005), and VCell (Moraru et al., 2008). Many of these tools are based on an interface that either allows a user to 'draw’ networks or enter information via a spreadsheet-like interface.

In addition to graphical tools, systems biology software developers have also created scripting languages that allow fine control over the kinds of simulations and analyses that are possible. One of the first general purpose systems biology scripting languages was Jarnac (Sauro and Fell, 2000), which implemented a Pascal-like syntax for describing models and simulations. More recently, general-purpose scripting languages such as Matlab, Python, Perl and R have gained widespread use within the scientific community. With the exception of Matlab, these languages are open source which makes them well suited for open research and reproducibility of computational experiments (Easterbrook, 2014).

Python has widespread use in academia as well as commercial settings due to its simple syntax, portability, and comprehensive standard library. These features make Python an ideal environment for computational and interactive work. In systems biology, there are a number Python software packages focused on modeling and simulation, including PySCeS (Olivier et al., 2005), SloppyCell (C.R. Myers et al., 2007), PySB (Lopez et al., 2013) and more recently Pycellerator (Shapiro and Mjolsness, 2015). These tools are generally focused on specific use cases and many require complex installation procedures or have limited user interfaces.

In this article we describe a new Python platform, Tellurium, designed to address these issues and add important new capabilities to the existing pool of Python-based Systems biology tools.

## 2 Design Rationale

Tellurium is based on a full, self-contained Python distribution which includes its own interpreter and is independent from any other Python distributions on the target machine. This avoids unnecessary interference with exiting Python installations. The included Python distribution is bundled with scientific packages and comes fully pre-configured, making it easy to copy and redeploy Tellurium to different machines. We utilize WinPython (https://winpython.github.io/) for distribution of the Python environment on Windows, whereas on Mac OS X we use the Apple disk image (.dmg) format. A Linux distribution is currently under development via conda (http://conda.pydata.org/).

We chose to use the Spyder IDE (https://github.com/spyder-ide/spyder) as the default platform for our software due to Spyder's straightforward and user-friendly interface akin to MATLAB (Figure 1A). Spyder's graphical user interface (GUI) is built with Python bindings for Qt (http://www.qt.io/), a stable and mature toolkit for cross-platform widget-based user interface development.

Tellurium comes with a wide variety of pre-installed Python libraries, plugins, and tools tailored for biological simulation, including packages to support common standards in systems and synthetic biology such as Systems Biology Markup Language (SBML) (Hucka et al., 2003), SED-ML (Waltemath et al., 2011), the COMBINE archive (Bergmann et al., 2014), and the synthetic biology design standard SBOL (Galdzicki et al., 2011; Galdzicki et al., 2014). SBML is a xml based *de facto* standard for representing cellular models. Similarly, SED-ML is a xml based standard for describing specific simulation experiments. The COMBINE archive is a zip based container for incorporating all information necessary for describing a model including but not limited to SBML, SED-ML files and data.

**Figure 1:**
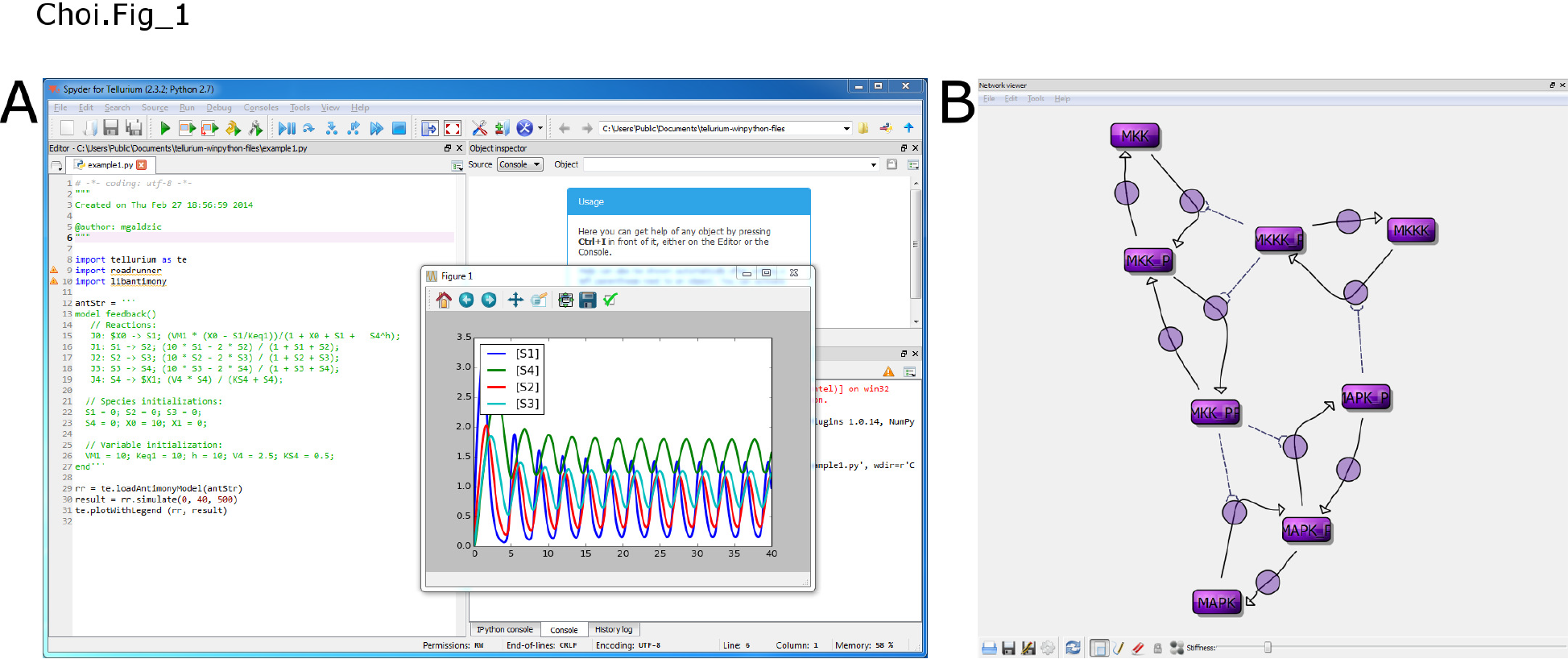
Screenshots from Tellurium. (A) Main interface of Tellurium (B) Output of network viewer plugin depicting MAPK cascade model.

## 3 Results

### 3.1 Libraries

#### Libraries Libroadrunner

Tellurium incorporates libRoadRunner (Somogyi et al., 2015) as the primary simulation engine. libRoadRunner is a high performance SBML simulation library which utilizes a custom Just-In-Time (JIT) compiler based on the LLVM compiler framework. libRoadRunner's solvers support ordinary differential equations (ODEs), discrete events, and stochastic timecourse simulations. libRoadRunner also supports steady state, sensitivity analysis and computes control coefficients and elasticities (Kacser and Burns, 1973; Heinrich and Rapoport, 1974). In the presence of conserved cycles, libRoadRunner can optionally use model reduction (Sauro and Ingalls, 2004; Vallabhajosyula et al., 2006).

Use of native machine code for simulation in libRoadRunner greatly increases the simulation speed, allowing users to perform simulations on large ensembles of models and multi-scale systems. libRoadRunner offers a range of numerical timecourse solvers, including CVODE from the SUNDIALs suite (Serban and Hindmarsh, 2005), RK4 (Press et al., 1988; Sauro, 2014) and RKF45 (Burden and Faires, 2011). libRoadRunner also includes a stochastic timecourse solver based on Gillespie's direct method (Gillespie, 1977; Sauro, 2012), which computes probabilistic trajectories of discrete reaction systems. All solvers expose solver properties, i.e. **rr.integrator.relative_tolerance**, if applicable.

#### Antimony

Tellurium uses Antimony (Smith et al., 2009) as a model definition language. Antimony is a human readable/writable language that can be used to define SBML-compatible models including hierarchical models. For example, a simple unimolecular reaction,

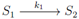

may be represented in Antimony as follows:

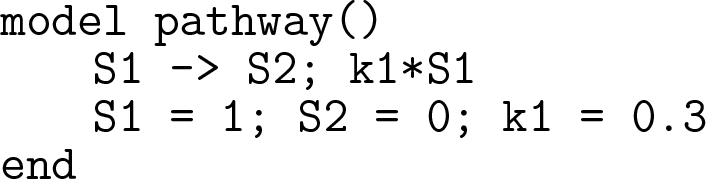

The second line describes the reaction and its kinetic law (*k*1·*S*1) while the third line initializes the reactant and product species (*S*1 and *S*2) and the rate constant *k*1. Antimony is designed to be a human-readable abstraction and an Antimony model definition can be converted to and from SBML. Tellurium provides helper methods that allow for direct construction of a **RoadRunner** instance from an Antimony string. Using the helper function **tellurium.loada()**, the previous example can be encoded and simulated in Python:

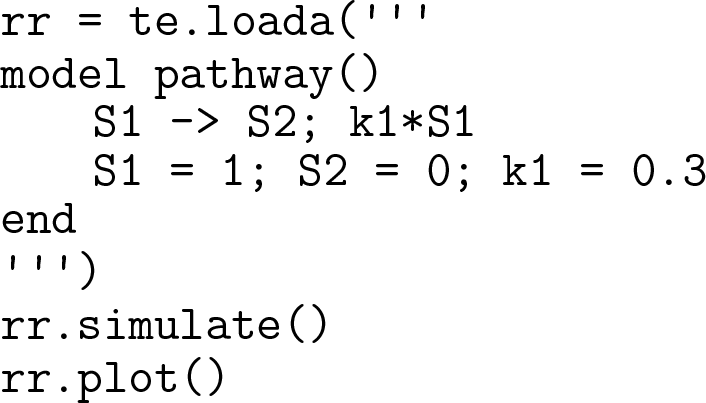

#### PhraSED-ML

PhraSED-ML (https://http://phrasedml.sf.net//) is a human-readable representation of SED-ML. By allowing users to specify simulation experiments in a human-readable form, we enable Tellurium to support reproducibility and exchangeability without sacrificing usability. The following example illustrates how a uniform timecourse simulation from t=0 to 100 with a 2D timecourse plot on species S1 and S2 can be described in phraSED-ML:

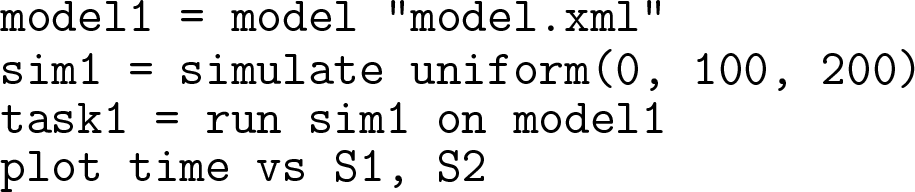

Tellurium supports direct execution of a phraSED-ML string through the **experiment()** module.

Tellurium comes with Python packages widely adopted in systems and synthetic biology community including libSBML (Bornstein et al., 2008), libSEDML (Waltemath et al., 2011; Bergmann et al., 2013), and pySBOL (https://github.com/SynBioDex/pySBOL). Common scientific libraries such as NumPy, SciPy, and matplotlib (https://www.scipy.org/) are included with Tellurium. Table 1 lists some of the pre-built libraries packaged with Tellurium.

**Table I:** Selected list of special-purpose systems / synthetic biology libraries and plugins included in Tellurium (beyond standard scientific Python packages such as NumPy and SciPy).

| **Packages** | **Spyder IDE Plugins** |
| --- | --- |
| libRoadRunner | Import SBML |
| Antimony | Import COMBINE archive |
| phraSED-ML | SBML network viewer |
| libSBML | Rate law templates |
| libSEDML |  |
| pySBOL |  |

### 3.2 IDE Plugins

One of the advantages of the Spyder IDE is that it has a modular plugin interface. This allows developers to customize the interface (Table 1).

#### Network Viewer

Tellurium includes a network viewer plugin which renders network diagrams encoded using the SBML Layout Extension (Gauges et al., 2006). The plugin is a complete rewrite of (Deckard et al., 2006) with cross-platform support and multiple language bindings. Internally, it utilizes SBNW (Medley et al., 2016), a C/C++ library for encoding and manipulating layout information in SBML files. For SBML files which lack layout information, the library uses the force-based Fruchterman-Reingold (FR) layout algorithm (Fruchterman and Reingold, 1991) to automatically generate and render network graphs containing SBML layout information. This layout information can then be written back to the file. Figure 1B shows a MAPK cascade model (Kholodenko, 2000) visualized using the network viewer plugin.

#### Import Plugin

The import plugin offers automatic conversion from SED-ML files and COMBINE archives to Python scripts and is designed to facilitate exchangeability and reproducibility of computational models. The plugin is invoked through a simple GUI and converts the input file or archive into an executable Python script. Although SED-ML files are designed to contain complete information on a particular simulation setup, the structure of scripts written in imperative languages differs inherently from that of a markup language with a specified data structure. Therefore, a SED-ML file must be translated at some point for Python to be able to execute the set of instructions supplied for the simulation. By providing a converter that automatically translates SED-ML files into Python code, we expect an increase in productivity while maintaining exchangeability of simulation setups through the use of a standardized format. Currently, our SED-ML to Python converter supports basic simulations (uniform timecourse, single step, and steady state convergence), repeated tasks, and three types of output (report, plot2D, and plot3D). In addition, our phraSED-ML notation serves as an intermediate representation which allows translating SED-ML files or combine archives to Tellurium-compatible scripts.

#### SBML Import

The SBML import plugin provides a simple and fast way to import and translate SBML models. SBML files can be directly loaded and immediately converted into embedded Antimony strings, ready to be used with any of Tellurium's simulation or analysis packages.

This is ideal for quickly checking the contents of an SBML file.

#### Rate law Template

The Rate Law Templates plugin provides a dictionary of commonly used rate equations in systems biology (Sauro, 2012). The plugin allows users to insert equations to the editor window. Currently, the library consists of 47 different rate laws which can be expanded through updating the rate law database.

### 3.3 Tellurium Libraries

Tellurium contains many Python packages that go beyond the capabilities of a standard scientific Python installation. These include parameter scanning, a straight-forward Python package for working with SBML, and various inter-format conversion tools. The supplementary material contains example scripts for each of the following features.

#### Parameter Scan Library

**teParameterScan** supplies functions to run multi-dimensional parameter scans. The module uses libRoadRunner to rapidly return the result of each step along the scan and provides several predefined functions for conveniently generating three dimensional and surface plots. The **teParameterScan** module also includes **teSteadyStateScan**, which is used for scanning through steady state values in linear or log space.

A simple example illustrating the use of **teParameterScan** is depicted in Figure 2. We used a model composed of said reactions:

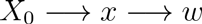

where species *X*_0_ and *w* are boundary species. The rate laws of the first and the second reactions are given by

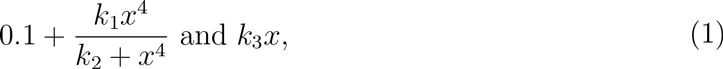

respectively. The initial concentration of species *x* has been changed, whose value varies from 0 to 1.8 in 10 steps. Figure 2 provides an explicit view of the timecourse concentration of *x* as the initial concentration changes.

**Figure 2:**
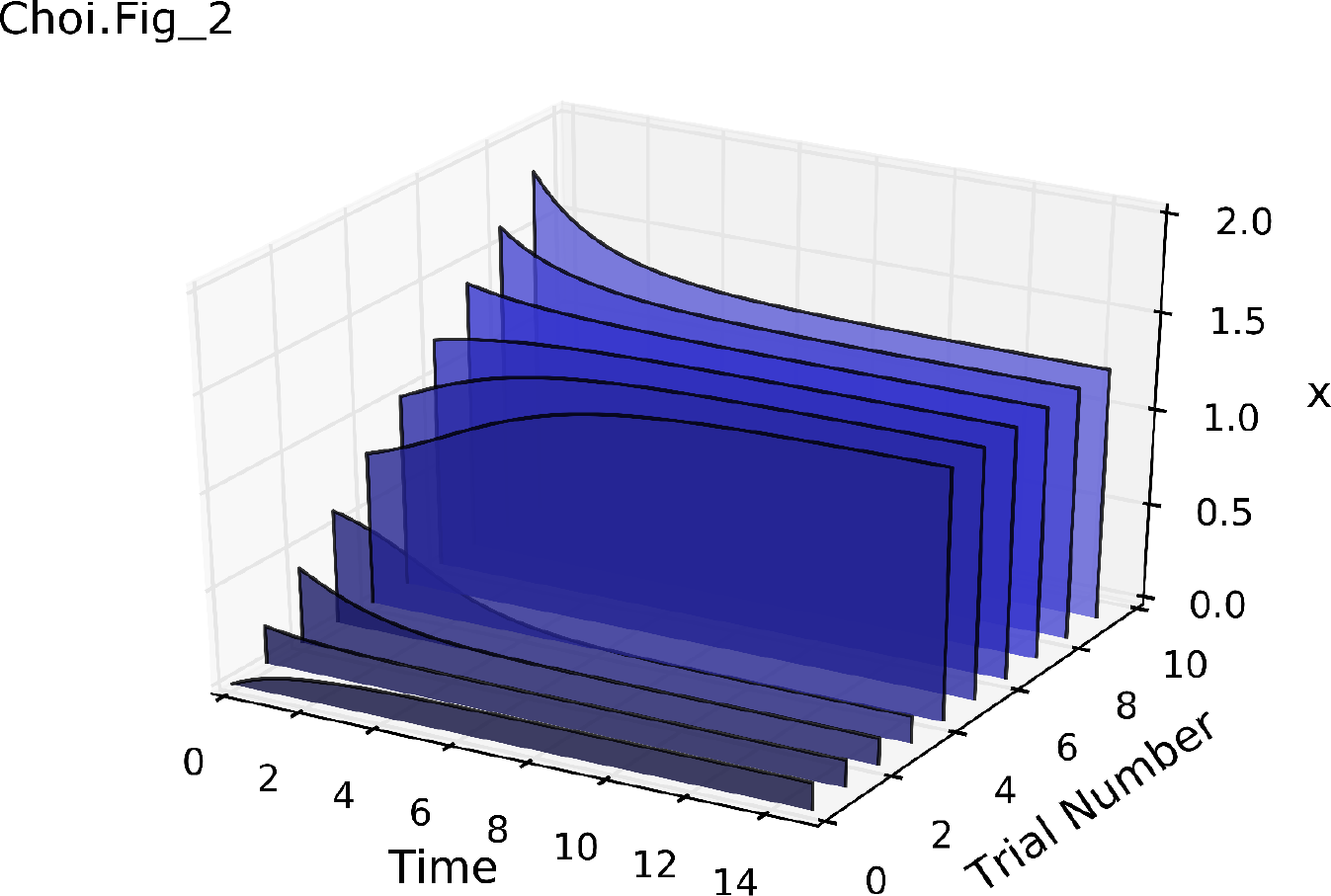
An example of **teParameterScan**. The figure depicts the result of a 1D parameter scan, with timecourses of species *x* concentration plotted for varying initial concentration of *x*.

#### Export to LaTeX: teExport

Support is provided to export plots in pgfplot format (Feuersänger, 2011). This is a standard plotting package for LaTeX. This package makes it simple to prepare publication quality plots. This is in addition to the usual PDF export provided by matplotlib.

#### SimpleSBML

SimpleSBML (Cannistra et al., 2015) allows users to generate SBML using simple Python scripts. It is intended for novice users who would rather not use the more complex libSBML Python API. SimpleSBML supports SBML Levels 2 and 3, and can be used to implement compartments, species, parameters, reactions, events, assignment rules, rate rules, and initial assignments.

#### MATLAB Support

MATLAB support is provided by the SBML2Matlab library (https://github.com/sys-bio/sbml2matlab). This library converts a standard SBML model into a MATLAB function which includes all the necessary information to run a timecourse simulation of the model from within MATLAB.

#### COMBINE Archive Support

Tellurium supports the COMBINE archive, a file format containing all the information necessary to reproduce a simulation experiment (Bergmann et al., 2014). Internally, Tellurium uses the **experiment** class to represent a simulation experiment, including the model (represented in Antimony), and the simulation setup (represented in phraSED-ML). These representations are automatically converted to SBML and SED-ML respective at export time. The result is a.omex or.zip file containing the SBML and SED-ML files as well as a manifest, as defined in (Bergmann et al., 2014).

### 3.4 Bifurcation Analysis

Bifurcation analysis enables qualitative changes in model behavior to be studied as a function of a model parameter. Such qualitative changes can include bistabiliy and oscillatory behavior (Angeli et al., 2004; Ermentrout and Terman, 2010). Tellurium supports bifurcation analysis through the AUTO2000 library (Doedel et al., 2002). The feature is implemented as a plugin for libRoadRunner, which allows it to directly access the simulation engine and perform computations without the overhead of a cross-language API. This also means that the bifurcation tool can be used outside of Python and hosted by other tools.

**Figure 3:**
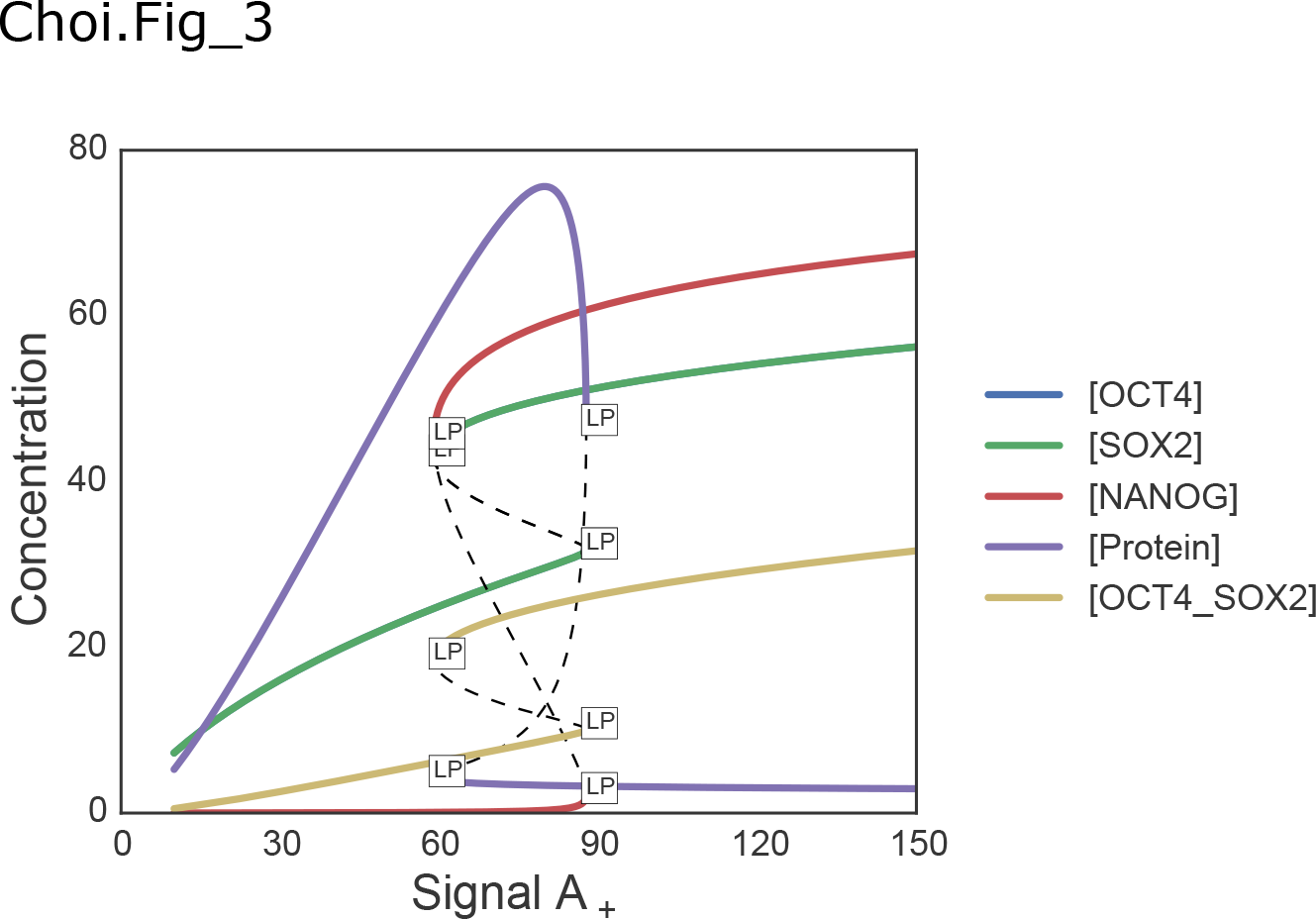
Bifurcation analysis applied to a model of an embryonic stem cell switch; Chickarmane et al., 2006. The label LP represents a fold or turning point bifurcation.

Tellurium's bifurcation plugin is designed to automatically compute a bifurcation in parameter space and plot a bifurcation diagram without user intervention. The user specifies a model parameter as the basis for the analysis. The plugin will then automatically scan a user-specified range of parameter values. If at some point the system changes to an alternate stationary state, the bifurcation is recorded and scanning continues. Figure 3 illustrates a number of bifurcation changes in a model of the embryonic stem cell switch (Chickarmane et al., 2006). For models where the stoichiometry matrix is not full rank (e.g. signaling pathways), libRoadRunner creates the appropriately reduced model to avoid numerical errors (Bergmann et al., 2006).

## 4 Applications

One of the strengths of Tellurium is its ability to take advantage of all the features of a scripting language that allows users to knit together the different libraries. The following applications illustrate how to use Tellurium's built-in solvers and tools in a typical systems biology workflow.

**Figure 4:**
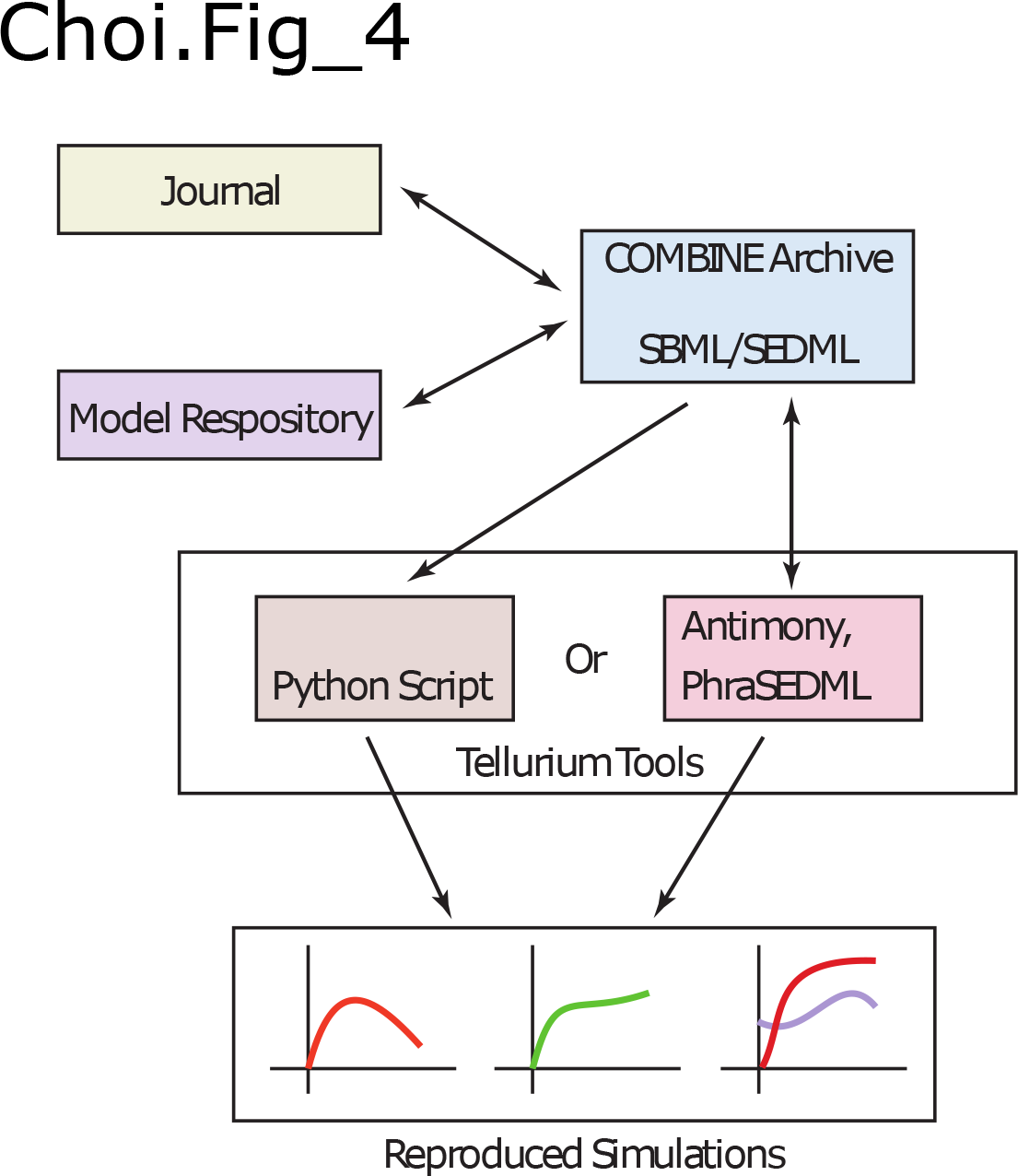
A COMBINE archive can be downloaded from a journal or model repository. Users can translate the COMBINE archive into either a Python or Antimony/phraSED-ML script. If the COMBINE archive provides sufficient information, users can reproduce each and every figure that the paper contains. If the archive is translated into an Antimony/phraSED-ML script, changes made to the model/simulation can be re-exported and submitted in the form of an updated archive to a journal, repository, or passed on to a colleague. This scenario therefore fully supports reproduction and exchange of computational models.

### 4.1 Reproducible Simulations using the COMBINE Archive

Support for reproducibility and exchangeability is one of the central features of Tellurium. To demonstrate this, we show how Tellurium can be used to create a COMBINE archive. In order to validate that the archive is written correctly, we use Frank Bergmann's suite of COMBINE archive utilities (https://github.com/fbergmann/CombineArchive). After validating the archive, we modify it and reload the result in Tellurium, demonstrating compliance with the COMBINE format and the complete cycle proposed in the workflow described in Figure 4.

**Figure 5:**
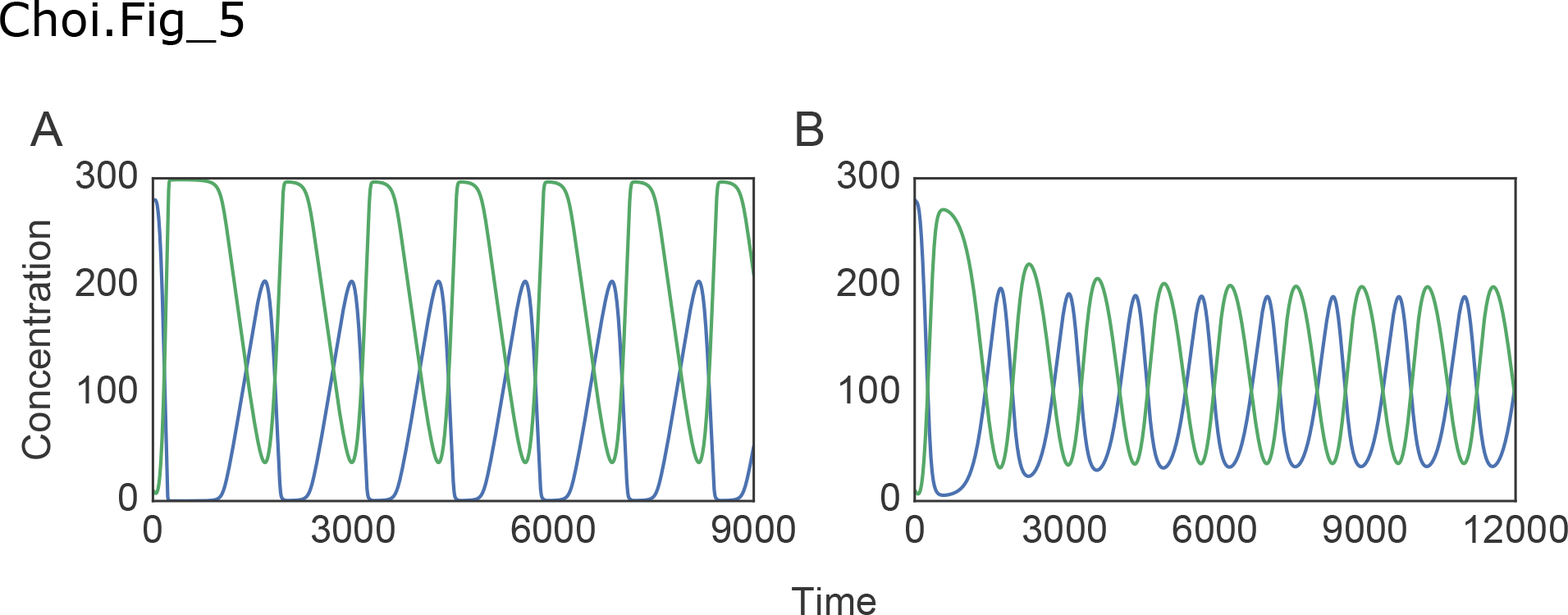
Timecourse simulation of MAPK and bi-phosphorylated MAPK concentration reproduced from (A) the original COMBINE archive, and (B) the modified combine archive. Blue lines represent the level of MAPK and green lines represent the level of bi-phosphorylated MAPK.

We start with the MAPK model described previously (Kholodenko, 2000). Using phraSED-ML, we define the simulation setup and the exact parameters used by the solver (see supplementary material for source code). The phraSED-ML representation is transparently translated into SED-ML (Figure 5A). Using the **exportAsCombine()** function provided in Tellurium's **teCombine** module, we then export the SBML and SED-ML components of the simulation to a COMBINE archive (.omex file).

Using the **CombineArchive** tool stated before, we can make direct changes to the SED-ML and SBML files embedded in the COMBINE archive. Here, we use the tool to obtain a different parameterization of the model that reproduces another figure in (Kholodenko, 2000). The changed parameters and values are listed in Supplementary Table S1.

Figure 5B illustrates the output of the re-imported model in Tellurium. The example illustrates how Tellurium's workflow incorporates automated handling of the exchange of reproducible simulations.

### 4.2 Monte Carlo Simulation of Linear Reaction Chain

Monte Carlo simulation is a method based on repeated random sampling of variables with given probability distributions. The example serves to illustrate the flexibility of using a scripting language.

**Figure 6:**
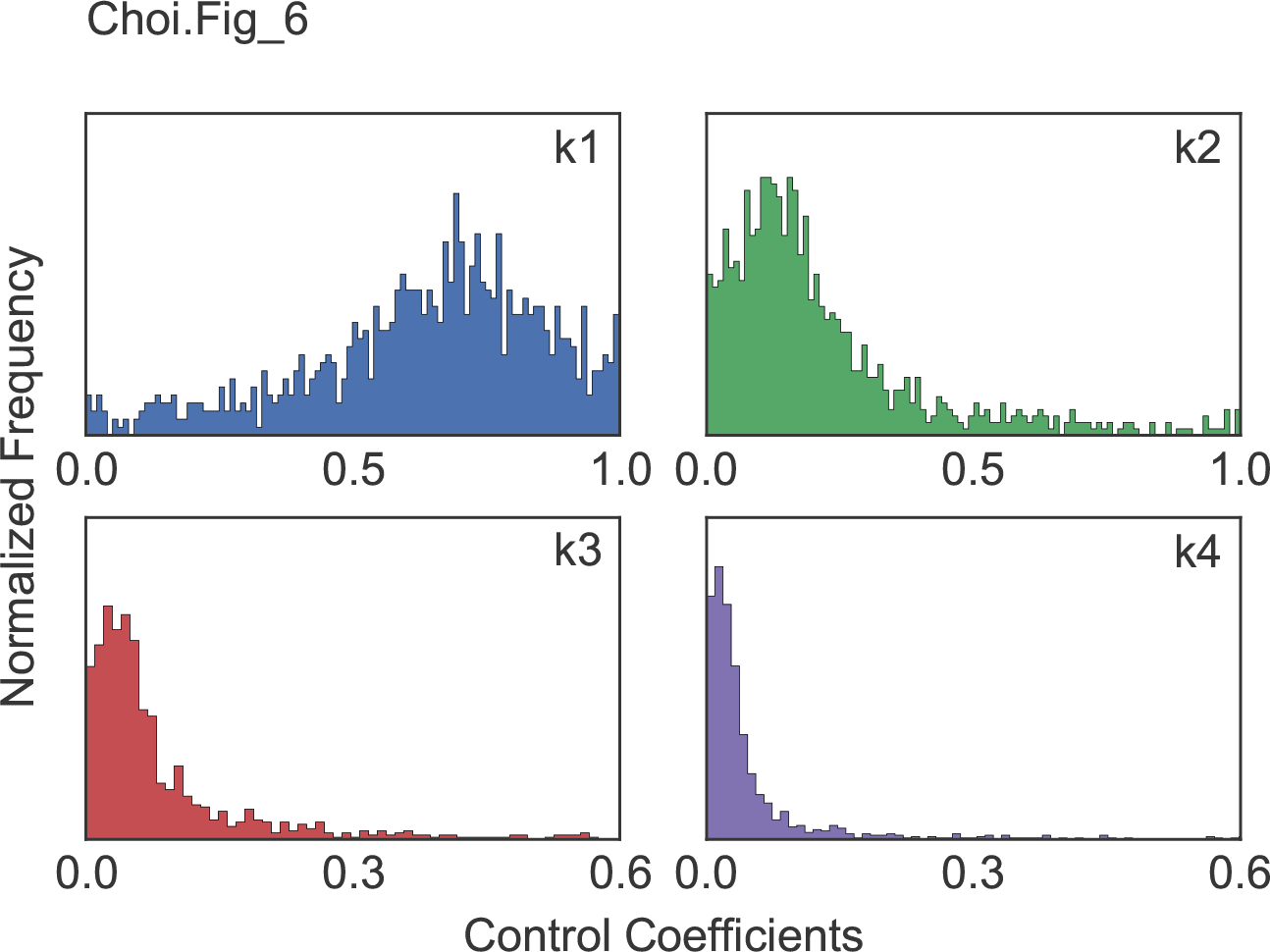
Normalized distribution of scaled flux control coefficients for four different rate constants with respect to the first reaction of a regular chain.

Here, we show that Tellurium can be used to visualize the distribution of flux control coefficients by performing a Monte Carlo simulation with rate constants as parameters. Consider a model of four reversible reactions, i.e.,

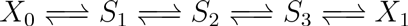

where *X*_0_ and *X*_1_ are boundary species and the rate law is defined by (Sauro, 2012)

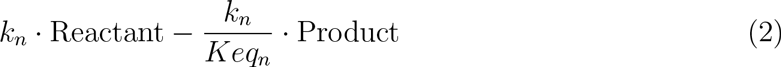

for each reaction *n*. For our example, we keep the value of *K_eq__n_* constant but randomly vary *k_n_.* The sampling is done 1000 times for each parameter *k*_n_ where a random parameter value is drawn from a uniform distribution with interval ranging from zero to ten. For each run, new parameter values are assigned to the roadrunner model and scaled flux control coefficients are calculated for each rate constant with respect to the first reaction of the regular chain. The sampled distributions of flux control coefficients are plotted in Figure6. The distribution reveal that the first rate constant, *k*_1_, has the largest effect on the first reaction with decreasing effects for the downstream reactions.

### 4.3 Parameter Confidence Intervals Estimation using Monte Carlo Bootstrapping

In parameter estimation, a confidence interval refers to the bounds on the estimated parameter value for a given likelihood. Bootstrapping estimates the effect of noise in the data on the fitted parameter values.

In this application, we show how to estimate the confidence intervals of fitted parameters through the use of the Monte Carlo bootstrap method (Sauro and Barrett, 1995; Sauro, 2014). Monte Carlo bootstrapping estimates confidence intervals by repeatedly resampling the experimental data used for the fitter. The standard procedure starts with initial estimation of parameters on original experimental data to obtain the residuals and the expected output calculated from the model using the estimated parameters. For each run of bootstrapping, a residual is randomly picked with replacement and added to the expected curve to create a unique synthetic data set, on which another parameter estimation is performed. This procedure is repeated multiple times to obtain a sample of fitted parameter values. This sample is then used to compute the confidence bounds. For a sample of parameter values, the 95% confidence interval is given by,

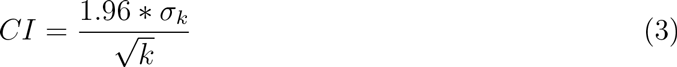

where *k* is the sampled values and σ is the standard deviation. We used the HIV protease model from (Kuzmič, 1996). This model contains the following set of reactions:

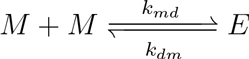

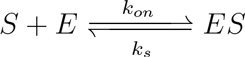

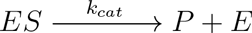

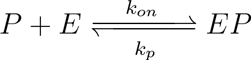

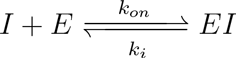

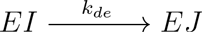

**Figure 7:**
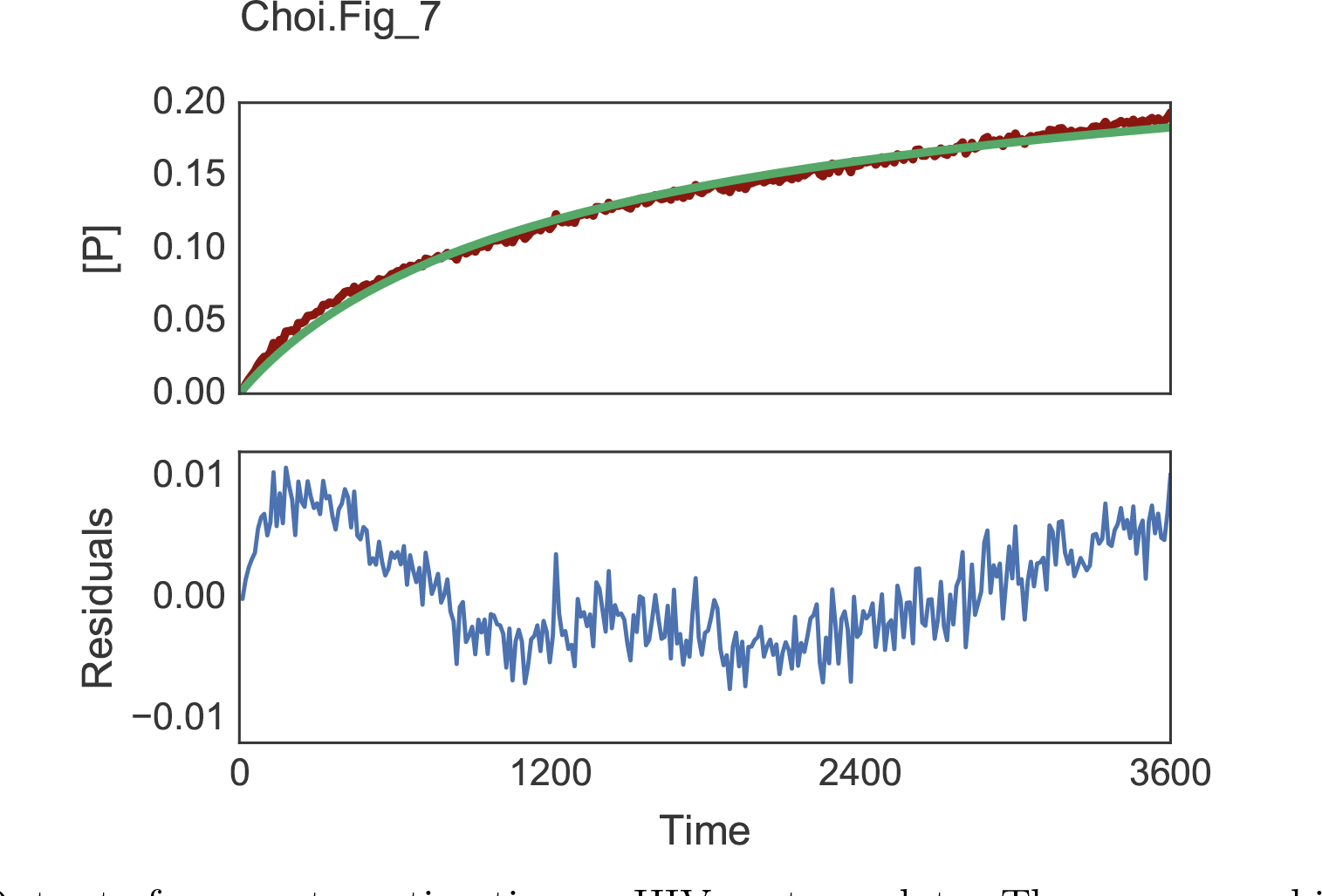
Output of parameter estimation on HIV protease data. The upper panel illustrates timecourse concentration of product P. The red line represents the raw data used for fitting and the green line represents timecourse data simulated through libRoadRunner using the fitted parameters. The lower panel shows the residuals between the raw and fitted line. Note the noticeable trend in the residual, which indicates an issue with the fitted model (Sauro, 2014).

We let *k_on_*=100, *k_dm_*=0.001, *k_md_*=0.1 and allowed the other five rate constants (*k_s_, k_cat_, k_p_, k_i_, k_de_*) to vary. We used libRoadRunner to perform timecourse simulations and compared the results to the experimental data set provided in (Kuzmič, 1996). For parameter estimation, we utilized the Nelder-Mead algorithm (Nelder and Mead, 1965) via the **lmfit** package (Newville et al., 2014) which provides simple wrapper functions for various optimization methods packaged with SciPy. The initial parameters are taken from (Kuzmič, 1996). Boundaries are set so that the parameter values do not become negative. Bootstrapping was repeated 500 times, which took about 5 minutes (302.4 ± 6.8 sec) on an Intel i7 4770 machine. Confidence intervals calculated from the Monte Carlo bootstrapping algorithm are listed in Supplementary Table S2. The upper panel in Figure 7 shows a typical outcome of a parameter estimation on the concentration of product *P*, where red dots represent experimental data and the green line represents the fitted curve. The residual of the fitting is illustrated in the panel below. It is also possible to check the correlation between fitted parameters, which is illustrated in Supplementary Figure S1.

## 5 Conclusion

Tellurium offers a powerful suite of modeling and simulation software tools in a user-friendly, accessible environment and can be used by researchers without specialized programming expertise.

Reproducibility and exchangability in biological models is a critical problem in the field of systems and synthetic biology. Several standards have been introduced to tackle the problem, but software support is crucial to enable adoption of these standards by end-users. Tellurium includes a number of tools which either allow the user to work directly with these standards, or transparently employ the standards when interchanging between software libraries.

We have endeavored to build a platform for collaboration by basing Tellurium on extensible and open architectures such as Spyder. Our tools are available under Open Source Initiative (OSI)-approved open source licenses. As a result, our users have the freedom to 6 apply additional customizations to Tellurium. Pervasive support for systems biology standards enables models created by Tellurium to be stored, reused, and modified reliably by other software tools.

## 6 Availability

Binary installers for Tellurium are available for Microsoft Windows and Mac OS X. Binaries, documentation, and full source code are available at http://tellurium.analogmachine.org. Tellurium is licensed under the Apache License Version 2.0. Linux distributions are being developed using conda.

## 7 Acknowledgments

This work was supported by the National Institute of General Medical Sciences of the National Institutes of Health under award numbers R01-GM081070 and the Federal Ministry of Education and Research (BMBF, Germany) within the research network Systems Medicine of the Liver (LiSyM) [grant number 031L0054]. The content is solely the responsibility of the authors and does not necessarily represent the official views of the National Institutes of Health.

